# Rhythmic expression of a cAMP phosphodiesterase switches neuronal excitability states in the mammalian suprachiasmatic nucleus

**DOI:** 10.64898/2026.07.14.738430

**Authors:** Priyam Narain, Marko Šušić, Sanzhar Kakenov, Giuseppe Antonio-Saldi, Justin Blau, Dipesh Chaudhury

**Affiliations:** Center for Genomics and Systems Biology, New York University Abu Dhabi, PO Box 129188, Abu Dhabi, UAE; Department of Biology, New York University Abu Dhabi, PO Box 129188, Abu Dhabi, UAE; Department of Biology, New York University, 24 Waverly Place, New York, NY 10003, USA; Center for Brain and Health, New York University Abu Dhabi, PO Box 129188, Abu Dhabi, UAE

## Abstract

How the transcriptome of a neuron determines its electrophysiological properties and how this change in different contexts are major questions in neurobiology. The intracellular molecular clocks in the circadian pacemaker neurons in the mammalian suprachiasmatic nucleus (SCN) drive 24hr rhythms in clock gene expression via a transcriptional-translational feedback loop^1,2^. These molecular clocks also control 24hr rhythms in the firing of SCN neurons^3–5^ – a subset of which enter a silent depolarized state around midday^6^. We used single-cell RNA sequencing and identified that the Avp⁺ / Nms⁺ and Vip⁺ / Nms⁺ are the SCN neurons that undergo daytime silencing. We also found high level and rhythmic expression of the *Pde10a* phosphodiesterase in these Nms⁺ SCN neurons. Acutely inhibiting Pde10a activity or elevating intracellular cAMP – to bypass high Pde10a levels – specifically shifts these Nms+ SCN neurons into a depolarized silent state hours ahead of schedule. A similar mechanism may occur in other neurons that switch to silent depolarized states.

## Introduction

Although we can determine the transcriptome of individual cells, it is still a challenge to understand how gene expression determines a cell’s functional properties, and how this can change in different contexts. This is particularly challenging in neurons, which select from a wide range of genes encoding ion channels and cell adhesion molecules to determine their firing properties and connections with other neurons – both of which often change over the lifetime of a neuron.

The mouse suprachiasmatic nucleus (SCN) contains approximately 20,000 neurons and is the principal circadian pacemaker in mammals that controls daily rhythms in physiology and behaviour^1,2,7^. Each SCN neuron contains a cell-autonomous molecular clock composed of core clock genes that operate through transcriptional–translational feedback loops (TTFLs), thereby sustaining rhythmic clock-gene expression^8,9^. These molecular clocks also control daily changes in the excitability and firing of SCN neurons that in turn drive circadian (24hr) rhythms in their outputs^3,5^.

The rhythmic excitability of SCN neurons is shaped by leak currents that set resting membrane potential and by voltage-gated Na⁺, Ca²⁺ and K⁺ channels that control firing. Daytime depolarization is supported by inward Na⁺ leak through NALCN, whereas nighttime hyperpolarization is promoted by outward K⁺ leak through TASK-3^10,11^. Other conductance further tunes this rhythm: persistent Na⁺ current and L-type Ca²⁺ current support daytime firing, whereas Kv12 and BK K⁺ currents contributing to reduced firing at night^12–14^. Importantly, daily changes in ionic current do not always match daily changes in channel mRNA. Some channels show rhythmic current without clear transcript rhythms, whereas others show transcript rhythms that do not predict the timing of the current. Thus, SCN excitability is shaped not only by channel expression, but also by post-transcriptional and post-translational regulation, including changes in channel gating, localization and phosphorylation.

Most SCN neurons fire at a higher frequency during the day than at night^3,15^. However, a subset of SCN neurons that express a *Per1-eGFP* transgene enter a highly depolarized but electrically silent state during the middle of the day, whereas the neighbouring eGFP^-^ neurons are less depolarized and continue to fire action potentials^6^.Silent depolarized states have also been described in cerebellar nuclei^16^, as well as in the lateral and medial habenula^17,18^. In addition, a subset of VIP-expressing SCN neurons remains active at night and contribute to anticipatory locomotor behaviour before dawn^19^.

Cyclic nucleotides also play an important role in the SCN. cAMP levels oscillate over 24 hours in the SCN^20^, and blocking cAMP production stops rhythmic clock gene expression^21^. SCN cAMP rhythms correlate inversely with Phosphodiesterase (PDE) activity^20^, suggesting that cyclic nucleotide degradation is under circadian regulation, but it is not known which PDE(s) are important in the SCN.

Thus, the genes that regulate the time-dependent neuronal physiological states in the SCN are poorly understood. In particular, little is known about how only a subset of SCN neurons enter a depolarized silent state. Here, we combined single-cell RNA sequencing with whole-cell patch-clamp recordings in *Per1-eGFP* and *Per1-Venus* reporter mice to define the molecular basis of rhythmic electrical states in SCN neurons. We confirmed that Per1-eGFP^+^ SCN neurons transition into a depolarized silent state during the middle of the day, and showed the same is true for Per1-Venus^+^ neurons. Single-cell profiling identified Avp^+^ / Nms^+^ and Vip^+^ / Nms^+^ neuronal populations as the principal reporter gene-positive SCN subsets. Our differential expression analysis revealed that expression of the *Pde10a* phosphodiesterase is enriched in GFP+ and Venus+ SCN neurons, and is also strongly rhythmic in those cells. We found that transiently inhibiting Pde10a or elevating intracellular cAMP selectively and rapidly shifted these Nms+ SCN neurons into a depolarized silent state, closely recapitulating their endogenous middle of the day phenotype. Thus, we conclude that temporal regulation of Pde10a-dependent cAMP signaling can switch a subset of SCN neurons between electrical states.

## Results

### Per1-eGFP⁺ and Per1-eGFP^-^ SCN neurons exhibit distinct electrophysiological states

Belle et al., 2009^6^ showed that Per1-eGFP+ SCN neurons enter a pronounced depolarized silent state and stop firing during the second half of the day. In contrast, they showed that Per1-eGFP-neurons maintain a less depolarized resting membrane potential (RMP) and continue firing action potentials until dusk.

We first wanted to confirm this heterogeneity in firing and so we performed targeted whole-cell patch-clamp recordings in acute SCN slices from *Per1-eGFP* transgenic mice. The data in Fig. 1 A-B shows that eGFP+ and eGFP-neurons exhibited similar RMP and spontaneous firing rates (SFR) between ZT2 and ZT5.75. However, the two populations showed distinct electrophysiological states between ZT6 and ZT11: eGFP+ neurons were ∼8 mV more depolarized than eGFP-neurons and stopped generating action potentials, whereas eGFP-neurons continued to fire at an average of ∼2 Hz.

**Figure 1.**
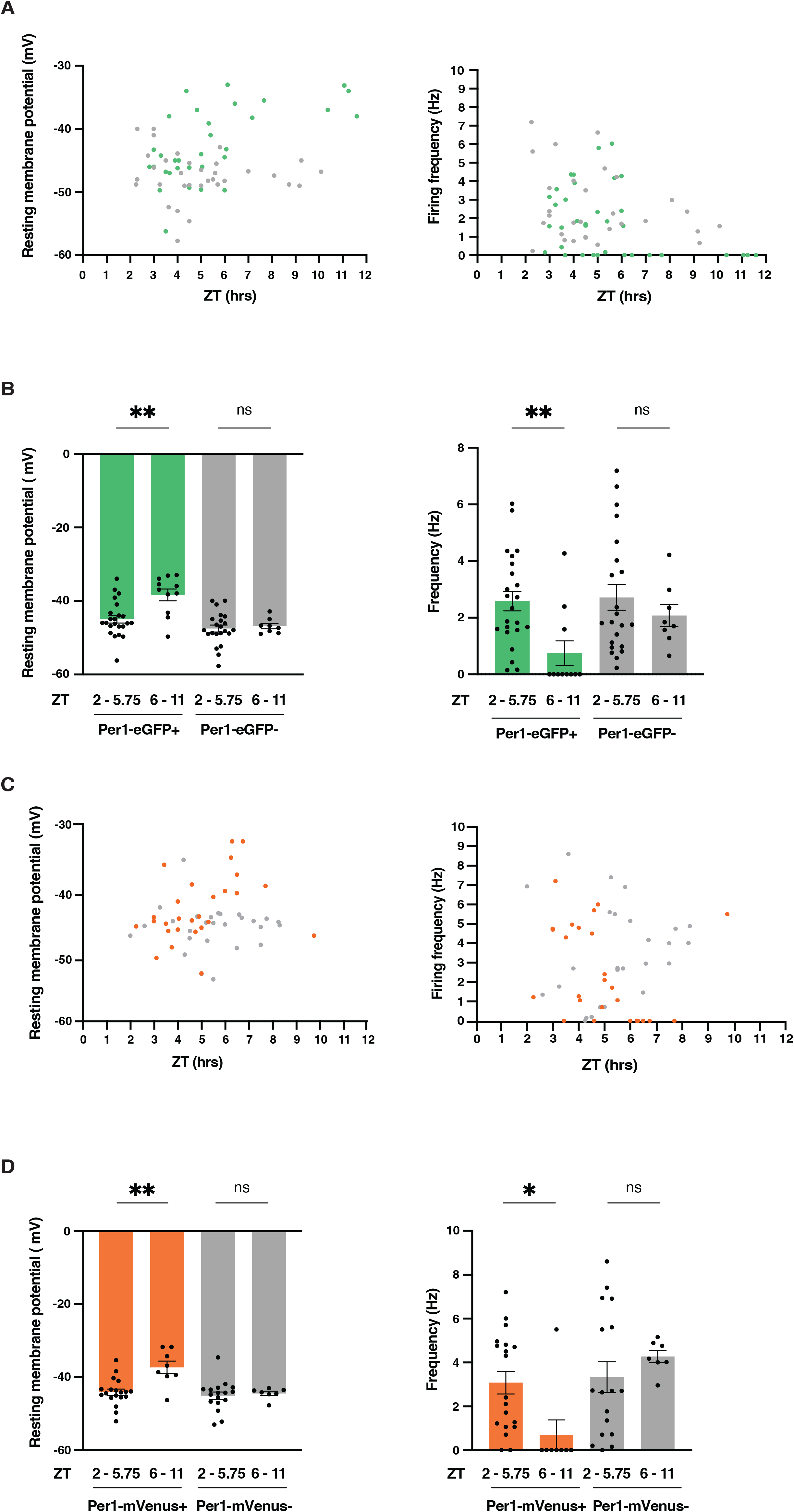
Electrophysiological properties of SCN neurons from *Per1-eGFP* and *Per1-Venus* mice. (A) Resting membrane potential (RMP mV, left) and spontaneous firing rate (SFR Hz, right) of individual Per1-eGFP⁺ neurons (green) and neighboring eGFP⁻ neurons (grey) recorded between ZT2 and ZT11 from acute coronal SCN slices prepared from 10-12 week old *Per1-eGFP* transgenic male mice. RMP was measured 1 min after establishing the whole-cell configuration. SFR was calculated from ≥ 1min of stable spontaneous firing. Each dot represents one neuron. (B) RMP (mV; left) and SFR (Hz; right) of Per1-eGFP⁺ and neighboring eGFP⁻ SCN neurons recorded at ZT2-5.75 and ZT6-11. In eGFP⁺ neurons, RMP shifted from −45.01 mV at ZT2–5.75 (n = 23) to −38.38 mV at ZT6–11 (n = 11; Welch’s unpaired two-tailed t-test, p = 0.0024). RMP did not change significantly in eGFP⁻ neurons over the same interval (−47.52 mV, n = 22, versus −46.85 mV, n = 8; p = 0.5773). SFR in eGFP⁺ neurons decreased from 2.59 Hz at ZT2–5.75 to 0.75 Hz at ZT6–11 (p = 0.0029), whereas SFR in eGFP⁻ neurons was not significantly altered (p = 0.2946). Bars show mean ± s.e.m.; each dot represents one neuron. p < 0.01; ns, not significant. (C) RMP (mV, left) and SFR (Hz, right) of individual Per1-Venus⁺ neurons (orange) and neighboring Venus⁻ neurons (grey) recorded between ZT2 and ZT11 from acute coronal SCN slices from 10-12 week-old *Per1-Venus* male mice. RMP was measured immediately after break-in. SFR was calculated from ≥ 1 min of stable spontaneous firing. Each dot represents one neuron. (D) RMP (mV, left) and SFR (Hz, right) of Venus⁺ and Venus⁻ SCN neurons recorded between ZT2-5.75 and ZT6-11. Venus⁺ neurons became significantly depolarized between ZT2-5.75 and ZT6-11, shifting from −44.08 mV (n = 19) to −37.34 mV (n = 8; Welch’s unpaired two-tailed t test, p = 0.0047). Venus⁻ neurons did not show a significant change in RMP over the same interval, measuring −45.03 mV at ZT2–5.75 (n = 17) and −44.41 mV at ZT6–11 (n = 7; p = 0.6003). Venus⁺ neurons also showed a significant reduction in SFR, from 3.075 Hz at ZT2-5.75 (n = 19) to 0.6875 Hz at ZT6-11 (n = 8; p = 0.0139). In contrast, SFR in Venus⁻ neurons was not significantly changed, measuring 3.329 Hz at ZT2–5.75 (n = 17) and 4.274 Hz at ZT6–11 (n = 7; p = 0.2232). Data are mean ± s.e.m. Statistical comparisons were performed using unpaired two-tailed t tests with Welch’s correction p < 0.05 = *, p < 0.01 = **, p < 0.001 = *** ; ns, not significant.

Depolarized silent states can indicate poor cellular health^6^. Therefore, we asked how these cells behave after injecting a steady hyperpolarizing current of up to -20 pA^6^ (see Methods). We found that this repolarized the RMP of eGFP⁺ neurons to an average of -46 mV (Supplementary Fig. 1A, left) and restored their ability to fire action potentials (Supplementary Fig. 1A, right). This confirms that the pronounced depolarized silent state of eGFP⁺ neurons is a physiological property of these cells at this time of day rather than a sign of poor health, as seen previously^6^.

### Per1-Venus⁺ and Per1-Venus^-^ SCN neurons also exhibit distinct electrophysiological states

To test if the electrophysiological properties of *Per1-eGFP*^+^ neurons are generalizable, we also performed whole-cell patch-clamp recordings in acute SCN slices from *Per1-Venus* mice (Fig. 1C-D). The *Per1-eGFP* transgene has a relatively short ∼3 kb fragment of the mouse *Per1* promoter driving destabilized GFP^22^, and probably does not fully recapitulate endogenous *mPer1* expression^23^.In contrast, the *Per1-Venus* reporter is a BAC transgene spanning the entire *mPer1* locus, including multiple regulatory regions that contribute to *Per1* expression, upstream of the rapidly maturing Venus-NLS-PEST reporter. *Per1-Venus* has slightly broader labeling in the SCN than *Per1-eGFP*^23,24^.

We found that the electrical properties of Venus⁺ and Venus^-^ SCN neurons were indistinguishable between ZT2 and ZT5.75. (Fig. 1C-D). However, differences in neuronal activity between Venus⁺ and Venus⁻ cells became evident between ZT6 and ZT11 (Fig. 1C-D): Venus⁺ neurons entered a depolarized silent state, whereas Venus⁻ neurons continued to fire with an average frequency of ∼4 Hz. We confirmed that the depolarized Venus⁺ neurons from were healthy as firing was restored after injecting a hyperpolarizing current (Supplementary Fig. 1B), as for Per1-eGFP+ SCN neurons^6^ (Supplementary Fig. 1A). Quiescent SCN neurons exhibiting a depolarized silent state have also been reported in rats^6^ and in the diurnal rodent *R. pumilio*^25^. Thus the transition to a depolarized, silent state seems to be a general property of a subset of SCN neurons.

### Single-cell profiling across the light–dark cycle defines SCN neuronal subtypes and eGFP⁺ populations

The rhythm of the SCN molecular clock determines the electrical properties of SCN neurons^4,5^. Given that SCN neurons show differences in the expression of *mPer1* reporter genes, it seemed likely that these electrophysiologically distinct SCN populations exhibit other differences in their transcriptomes. To test this, we performed single-cell RNA (scRNA) sequencing of SCN tissue from *Per1-eGFP* mice collected at six time points across a 12 hour: 12-hour light: dark (LD) cycle. Using unsupervised clustering and dimensionality reduction optimized to avoid over-clustering (UMAP; resolution = 0.6), we identified 24 distinct neuronal and non-neuronal clusters across all cells and time points, which segregated clearly in low dimensional space (Fig. 2A). The neuronal clusters identified in our datasets closely match recent SCN transcriptomic studies^19,26–29^ (Fig. 2B).

**Figure 2.**
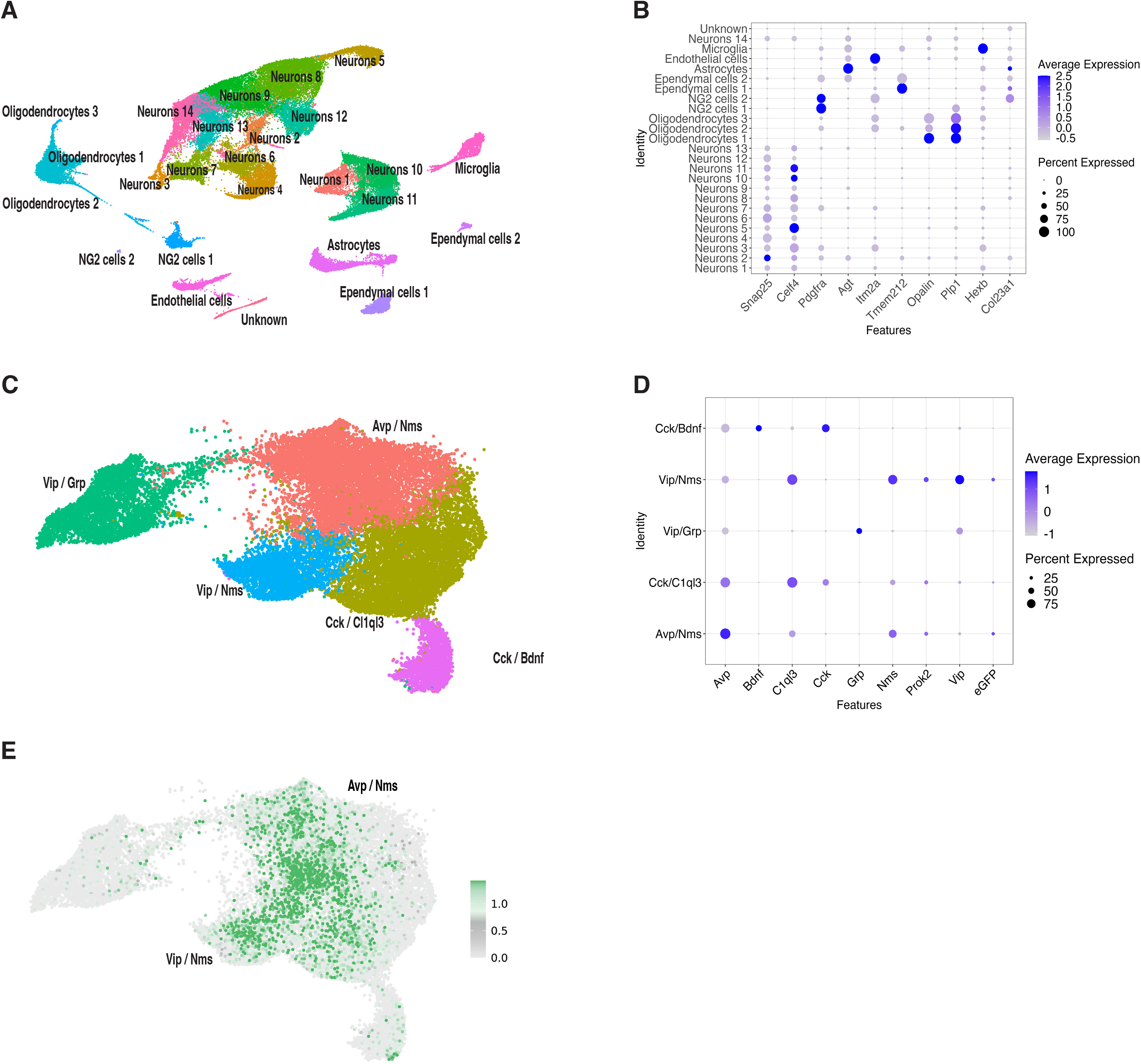
Single-cell RNA sequencing of SCN from *Per1-eGFP* mice identifies major cell classes and neuronal subtypes. (A) Integrated UMAP projection of single-cell transcriptomes from SCN cells collected at timepoints from ZT2 to ZT22. Each point represents one cell and is colored by its major cell class, assigned based on marker gene expression (see Methods). Identified classes include neurons, astrocytes, microglia, oligodendrocytes, ependymal cells, endothelial cells, and NG2 oligodendrocyte precursors. An additional Unknown population was detected but was excluded from further analysis. (B) Dot plot showing expression of canonical marker genes across SCN cells. Rows indicate transcriptionally defined cell identities and columns show representative marker genes. Dot size reflects the percentage of cells within each cluster expressing a gene, and color intensity indicates scaled average expression. Neuronal clusters are all enriched for *Snap25* and *Celf4* expression. NG2 cells express *Pdgfra*, oligodendrocytes show strong *Plp1* expression, and microglia are marked by *Hexb*. (C) UMAP projection of SCN neuronal subclusters colored by cluster identity. SCN-specific neuronal populations were defined by Scg2 expression together with established SCN neuropeptide markers. Sub-clustering was performed at a resolution of 0.2 to prevent over clustering. Two transcriptionally undefined clusters were excluded. Five SCN neuronal clusters were retained for downstream analysis. (D) Dot plot showing expression of canonical SCN neuropeptide markers across neuronal subclusters. Rows represent transcriptionally defined neuronal identities (Avp / Nms, Cck / C1ql3, Vip / Grp, Vip / Nms, and Cck / Bdnf), and columns represent selected marker genes (Avp, Bdnf, C1ql3, Cck, Grp, Nms, Prok2, Vip, and eGFP). Dot size indicates the percentage of cells expressing each gene within a cluster. Color intensity reflects average expression from low (pale) to high (dark). (E) UMAP projection of neuronal subclusters defined in C, colored by normalized eGFP expression. Each point represents a single cell, with color intensity indicating relative eGFP abundance (grey, low; green, high). eGFP expression is enriched in Avp / Nms and Vip / Nms clusters.

For further analysis, we selected neuronal populations showing high expression of previously published marker genes^26,28^, including *Scg2* (Secretogranin II), *Rgs16* (Regulator of G protein signaling 16), *Lhx1* (LIM homeobox 1), and several neuropeptides (*Avp, Vip, Nms, Grp, Cck, Bdnf, C1ql3*) (Supplementary Fig. 2). We defined 5 main neuronal populations (Fig. 2C, D). Two clusters expressed Neuromedin S (*Nms*), and co-expressed either Arginine Vasopressin (Avp⁺ / Nms⁺) or Vasoactive Intestinal Peptide (Vip⁺ / Nms⁺). Two clusters also expressed Cholecystokinin (Cck) and co-expressed either Brain-Derived Neurotrophic Factor (Cck^+^/ Bdnf^+^) or Complement C1q-like protein 3 (Cck^+^/ C1ql3^+^). We also identified a Vip⁺ / Gastrin Releasing Peptide (Vip^+^ / Grp^+^) neuronal cluster. We excluded two smaller clusters from downstream analyses as they lacked clear enrichment for the canonical neuropeptide SCN markers^26^.

Marker analysis using Seurat demonstrated that GFP expression is a defining marker for the two Nms+ populations (Fig. 2E). GFP expression was also detected at low levels in the Cck⁺/C1ql3⁺ cluster in the dot plot (Fig. 2D), but GFP was not defined as a cluster marker. Based on these annotations, we grouped the Avp⁺/ Nms⁺ and the Vip⁺/ Nms⁺ clusters together as GFP^+^ neurons, and the Vip⁺ / Grp⁺, Cck⁺ / C1ql3⁺, and Cck⁺ / Bdnf⁺ subclusters as GFP^-^neurons for downstream analysis.

### Clock-gene rhythms differ mainly in phase across SCN neuronal subtypes

We first examined whether the major SCN neuronal subtypes showed the expected rhythms in clock gene expression. Wen et al. reported that rhythmic clock gene expression is most prominent in *Avp*⁺/*Nms*⁺, *Vip*⁺/*Nms*⁺ and *Cck*⁺/*C1ql3*⁺ neurons, whereas *Grp*⁺/*Vip*⁺ and *Cck*⁺/*Bdnf*⁺ neurons show weaker rhythmicity^26^. They also found subtype-dependent phase differences, including a phase shift of *Cck*⁺/*C1ql3*⁺ neurons relative to *Avp*⁺/*Nms*⁺ and *Vip*⁺/*Nms*⁺ neurons. In our LD dataset (Figure 3A), we made similar findings. *Per1*, *Per2* and *Arntl* showed clear rhythmic expression in *Avp*⁺/*Nms*⁺, *Vip*⁺/*Nms*⁺ and *Cck*⁺/*C1ql3*⁺ neurons, with the expected temporal order: *Per1* peaked during the early day, *Per2* peaked later, and *Arntl* peaked around ZT14.*Cck*⁺/*Bdnf*⁺ neurons also showed rhythmic expression of some core clock genes, particularly *Per1* and *Per2*, whereas *Vip*⁺/*Grp*⁺ neurons showed weaker clock gene rhythmicity. Thus, our dataset reproduces the major subtype-specific clock gene patterns reported previously, while extending them to Per1-eGFP+ neurons in LD.

**Figure 3.**
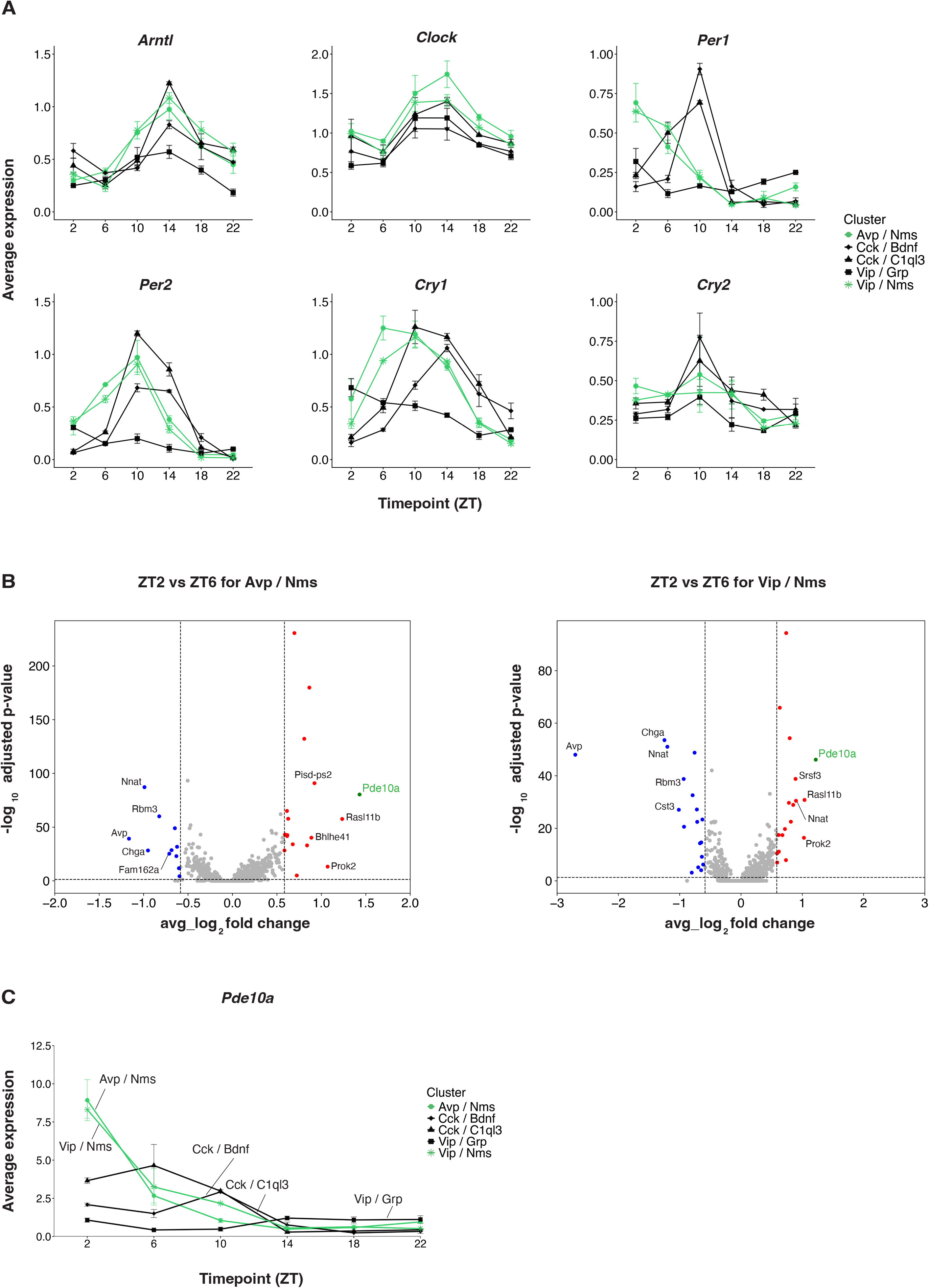
Expression of clock genes and *Pde10a* in SCN neuronal clusters. (A) Temporal expression of clock genes in SCN neuronal sub clusters. Line plots showing the average expression levels of six core clock genes (*Arntl*, *Cloc*k, *Per1*, *Per2*, *Cry1* and *Cry2*) across a 24-hour LD cycle. Each panel represents a single clock gene, with individual lines denoting the average expression profile in each of the 5 major SCN neuronal subclusters: Avp^+^ / Nms^+^ (green), Cck ^+^/ C1ql3^+^ (black), Vip^+^ / Grp^+^ (black), Vip^+^ / Nms^+^ (green), and Cck^+^ / Bdnf^+^ (black). (B) Differential gene expression analysis between ZT2 and ZT6 in eGFP⁺ SCN neurons. Left: Avp⁺/Nms⁺ eGFP⁺ neurons. Right: Vip⁺/Nms⁺ eGFP⁺ neurons. The x-axis shows log₂ fold change (ZT2 relative to ZT6), and the y-axis shows −log₁₀ adjusted p value. Genes enriched at ZT2 are shown in red, whereas genes enriched at ZT6 are shown in blue. Non-significant genes are shown in grey. Selected genes are annotated, including *Pde10a* (shown in green), which is upregulated at ZT2 in both neuronal populations. Differential expression was assessed using a Wilcoxon rank-sum test (adjusted p ≤ 0.05, fold change ≥ 1.40). (C) Average Pde10a RNA levels over time in the five transcriptionally defined SCN neuronal subtypes. *Pde10a* was rhythmically expressed in the reporter-enriched Avp⁺/Nms⁺/eGFP⁺ and Vip⁺/Nms⁺/eGFP⁺ populations, where expression was highest during the early day and declined at later time points (Avp⁺/Nms⁺, q = 0.0038; Vip⁺/Nms⁺, q = 0.021). *Pde10a* was also rhythmic in Cck⁺/C1ql3⁺ and Cck⁺/Bdnf⁺ neurons, but with later temporal profiles (Cck⁺/C1ql3⁺, q = 9.95 × 10⁻⁶; Cck⁺/Bdnf⁺, q = 3.28 × 10⁻⁴). In contrast, Vip⁺/Grp⁺ neurons showed low expression and no significant rhythmicity (q = 0.268). q values are Benjamini–Hochberg-adjusted rhythmicity p values.

### *Pde10a* is temporally regulated in eGFP⁺ SCN neuronal subtypes

We next compared gene expression between ZT2 and ZT6 to link the transcriptional state of SCN neurons to their physiology. We chose these times because ZT2 is when eGFP⁺ SCN neurons are active, and ZT6 is when they enter the depolarized silent state (Fig. 1). We performed differential expression analysis separately for Avp⁺/ Nms⁺/ eGFP⁺ and Vip⁺/ Nms⁺/ eGFP⁺ neuronal populations using a Wilcoxon rank-sum test. We identified 44 differentially expressed genes in Avp⁺ / Nms⁺ eGFP⁺ neurons and 45 differentially expressed genes in Vip⁺ / Nms⁺ eGFP⁺ neurons applying thresholds of 1.40 for fold change, and adjusted p ≤0.05, (Fig. 3B).

Of these differentially regulated genes with p ≤0.05, 228 genes are shared between the Avp⁺ / Nms⁺ eGFP⁺ and Vip⁺ / Nms⁺ eGFP⁺ SCN neurons. We did not observe any ion channel genes in this list. *Phosphodiesterase 10a* (*Pde10a*) is the most strongly up-regulated gene at ZT2 vs ZT6 in both clusters (Fig. 3B): *Pde10a* RNA levels were 2.7-fold higher in Avp⁺ / Nms⁺ neurons at ZT2 than ZT6, and 2.3-fold higher in Vip⁺ / Nms⁺ neurons.

Because *Pde10a* showed higher expression at ZT2 in eGFP⁺ Avp⁺/Nms⁺ and Vip⁺/Nms⁺ neurons, we next examined its rhythmicity across the day (Fig. 3C). *Pde10a* was rhythmic in multiple SCN neuronal subtypes, but its phase differed between clusters. In the eGFP^+^ populations, *Pde10a* peaked during the early day and declined at later ZTs, whereas eGFP⁻ *Cck*⁺ populations showed distinct temporal profiles and *Vip*⁺/*Grp*⁺ neurons showed weaker rhythmicity. Thus, eGFP⁺ neurons are not defined by exclusive *Pde10a* expression, but by a shared early-day peak in Pde10a. This timing may be relevant to their electrophysiological rhythm, as these same neurons later transition into a depolarized low-firing state. Importantly, *Pde10a* was 18.24-fold higher at ZT2 than ZT14 in Avp⁺/Nms⁺/eGFP⁺ neurons and 15.18-fold-fold higher in Vip⁺/Nms⁺/eGFP⁺ neurons.

### Transient inhibition of Pde10a selectively shifts eGFP⁺ neurons to a depolarized silent state

To test whether Pde10a contributes to the electrophysiological state of eGFP⁺ SCN neurons, we used papaverine (Ppv) to acutely inhibit Pde10a activity in SCN slices. Papaverine has been widely used in slice and in vivo studies to inhibit Pde10a^30–32^. Although papaverine is not fully selective for Pde10a^33^ the other 23 Pde genes examined showed low or nearly undetectable expression (Supplementary Fig. 3). We performed whole-cell current-clamp recordings between ZT2 and ZT5.75 when both eGFP⁺ and eGFP⁻ neurons are spontaneously active. We identified eGFP^+^ and eGFP^-^ neurons by their fluorescence levels, and recorded baseline RMP and SFR prior to bath applying 50 μM papaverine. The data in Fig. 4A-B show that papaverine depolarized the RMP of eGFP⁺ neurons by an average of 8 mV (left), and either reduced firing frequency or completely silenced these cells (right), with effects typically detected after ∼4 mins of application. In contrast, eGFP⁻ neurons showed minimal changes in RMP (Fig. 4A-B) and firing frequency (Fig. 4A-B).

**Figure 4.**
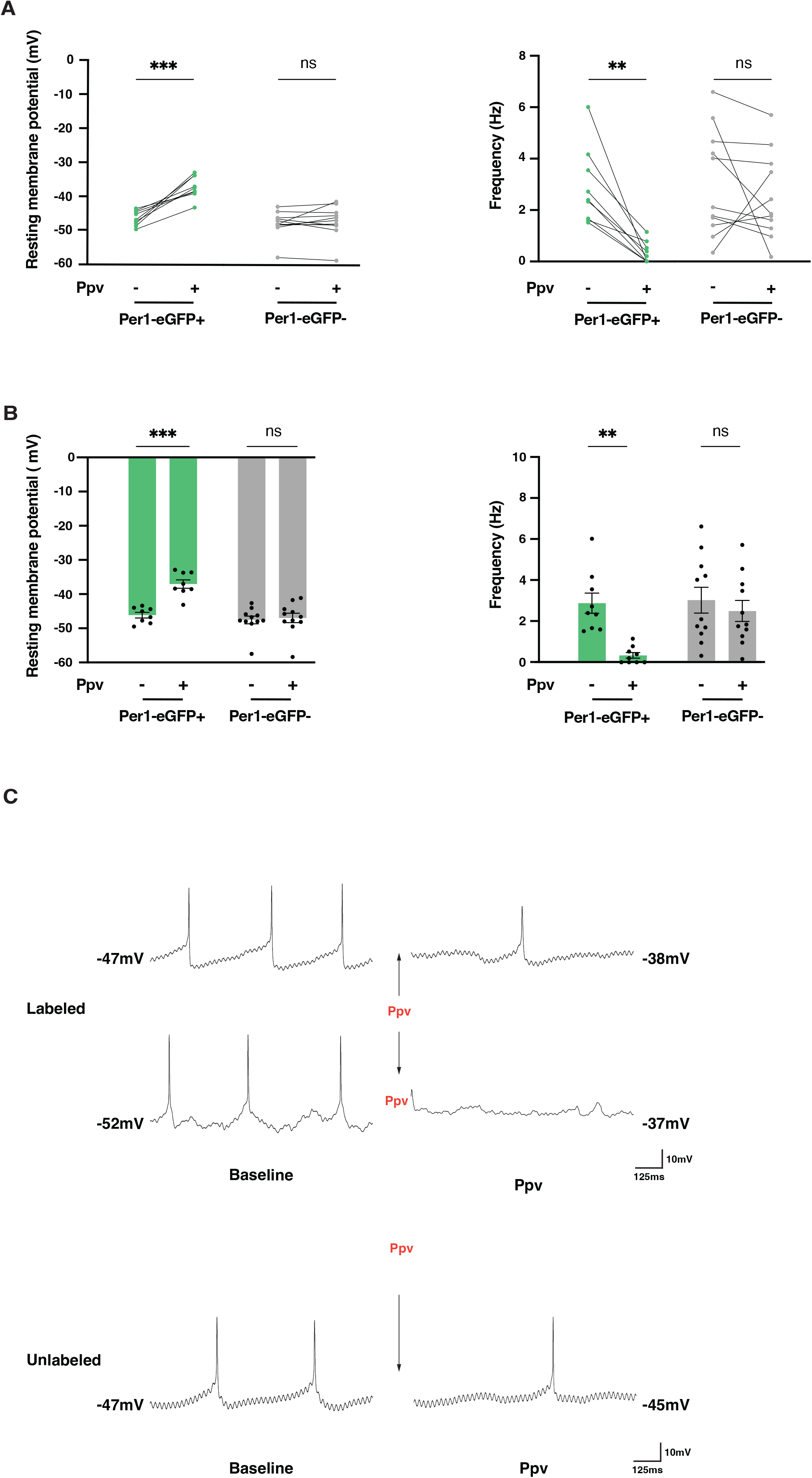
Papaverine depolarizes and silences eGFP⁺ SCN neurons but not eGFP⁻ neurons. Electrophysiological recordings were obtained from SCN neurons between ZT2 and 5.75, before (−) and after (+) bath application of the Pde10a inhibitor papaverine (Ppv). (A) RMP (left) and SFR (right) of eGFP⁺ and eGFP⁻ neurons before (−) and after (+) adding 50 µM papaverine. Adding papaverine depolarized eGFP⁺ neurons (paired two-tailed t-test, p = 0.0001, n = 9) but did not affect eGFP⁻ neurons (p = 0.4481, n = 11). Adding papaverine reduced firing in eGFP⁺ neurons (p = 0.0010) but not in eGFP⁻ neurons (p = 0.4326). Each line represents one neuron. (B) Quantification of RMP (left) and SFR (right) in eGFP⁺ and eGFP⁻ neurons before (−) and after (+) adding 50 µM papaverine. Bars represent mean ± s.e.m., and dots indicate individual neurons. **p < 0.01; RMP: ***p < 0.001. (C) Representative current-clamp recordings from eGFP⁺ and eGFP⁻ neurons before and after application of 50 µM papaverine. The top two examples are from eGFP⁺ neurons and show depolarization with reduced action potential firing after papaverine. The bottom example is from an eGFP⁻ neuron and shows little change after papaverine. Baseline traces were recorded 1 min after establishing the whole-cell configuration, and papaverine traces represent effect after 4 min of drug perfusion. Scale bars: 10 mV, 125 ms.

In the examples shown (Figure. 4C), eGFP⁺ neurons depolarized from approximately −47 to −38 mV and from −52 to −37 mV after adding papaverine, with a marked reduction in action potential firing. In contrast, the eGFP-neuron in Figure. 4C showed little change in membrane potential or firing pattern after papaverine. To exclude patch instability or compromised cell health as a cause of silencing, eGFP⁺ neurons were hyperpolarized after papaverine treatment. These cells remained capable of firing upon current injection, indicating they are healthy.

Thus, applying papaverine early in the day selectively shifts eGFP⁺ neurons into a depolarized, low-firing state, resembling that normally observed for eGFP⁺ neurons in the second half of the day (Fig. 1A, B). Although we cannot exclude a contribution from other PDEs, our expression data points to *Pde10a* RNA levels as highest in the SCN (Supplementary Fig. 3). Thus, the effect of papaverine is consistent with inhibition of Pde10A in these cells. The effect of papaverine is also consistent with *Pde10a* expression normally decreasing in eGFP⁺ neurons between ZT2 and ZT6 to permit the depolarized silent state by ZT6. This predicts that Pde10a is a fairly short-lived protein and thus that Pde10a protein levels closely follow *Pde10a* RNA levels.

### *Pde10a* is rhythmically expressed in Per1-Venus⁺ SCN neurons

Given that SCN neurons in *Per1-Venus* mice exhibit electrophysiological properties similar to those in *Per1-eGFP* mice (Fig. 1), we asked whether they share the same differentially expressed genes. We therefore performed scRNA sequencing of SCN tissue from *Per1-Venus* mice (Fig. 5A). This time we only isolated the SCN from mice at ZT2, ZT6, ZT10 and ZT14 because we were most interested in genes that could explain the different electrophysiological properties of SCN neurons before and after ZT6 (Fig. 1C, D).

**Figure 5.**
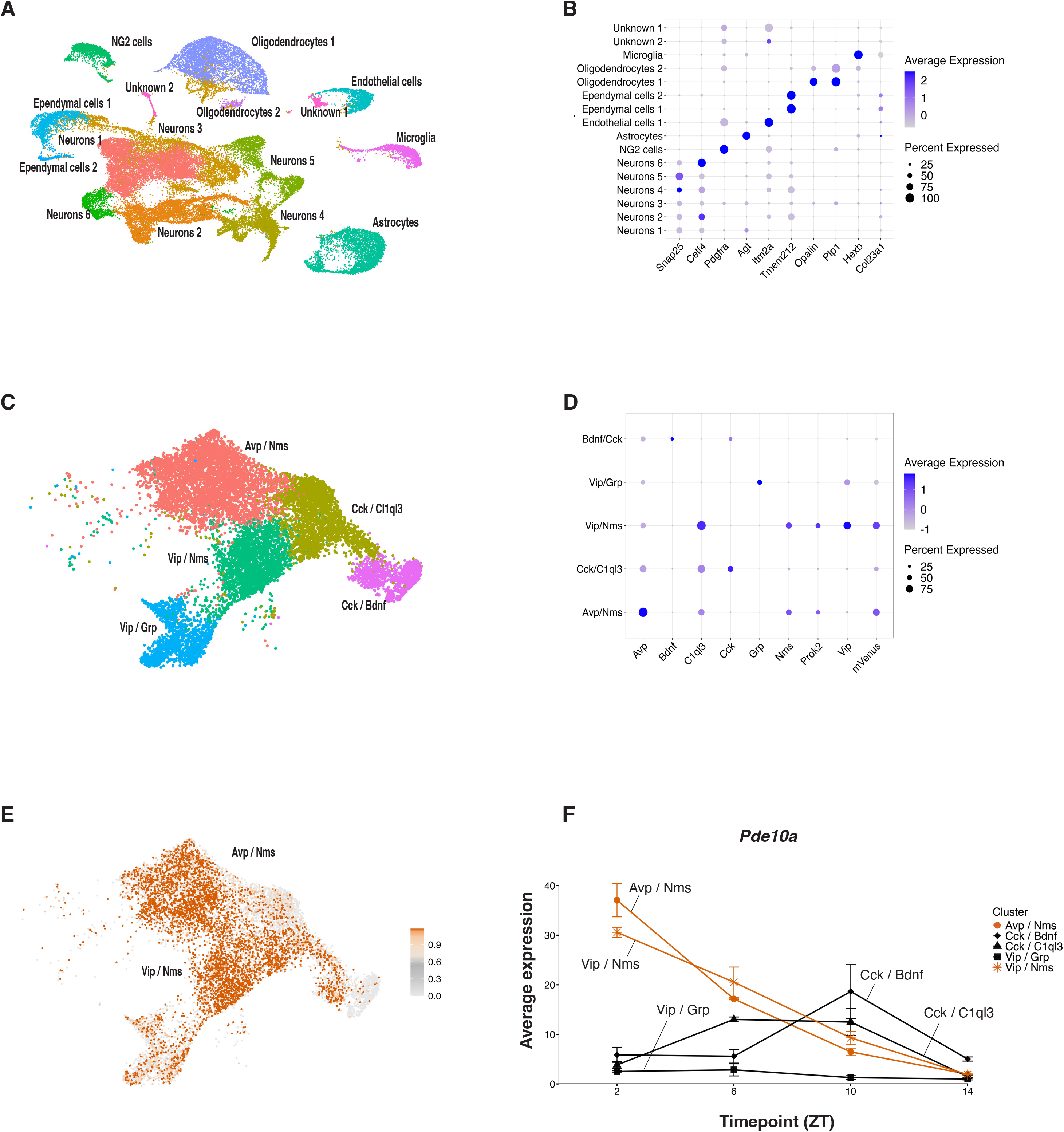
Single-cell transcriptomics reveals neuronal subtypes and rhythmic *Pde10*a expression in the SCN of *Per1-Venus* mice. (A) UMAP projection of single-cell transcriptomes from SCN tissue collected at four times in LD from *Per1-Venus* mice (ZT2, ZT6, ZT10 and ZT14). Each point represents one cell, colored by annotated cell identity, including neuronal and non-neuronal populations. (B) Dot plot showing expression of canonical marker genes across SCN cell classes. Dot size indicates the percentage of cells expressing each gene. Color intensity represents scaled average expression of that gene. (C) UMAP projection of reclustered SCN neurons identifying five neuronal subtypes defined by neuropeptide expression. (D) Dot plot showing expression of neuropeptides and Venus in the 5 different subtypes of SCN neurons. Venus expression is highest in Avp+ / Nms+ and Vip+ / Nms+ neurons. (E) UMAP showing distribution of *Venus* expression from low (grey) to high (orange) superimposed on the cluster plot from C. (F) Average Pde10a RNA expression at ZT2-ZT14 in the five transcriptionally defined SCN neuronal subtypes from Per1-Venus mice. In the Venus expressing Avp⁺/Nms⁺ and Vip⁺/Nms⁺ populations, *Pde10a* expression was highest at ZT2 and lowest at ZT14, with ZT2/ZT14 expression ratios of 18.42 and 17.83, respectively. Venus⁻ populations also expressed Pde10a but showed lower early-day expression and different temporal profiles.

SCN neurons separated into clusters defined by the same neuropeptide markers as in Per1-eGFP mice (Fig. 5B, C, Supplementary Fig. 4). Venus was identified as a marker gene for the Avp⁺ / Nms⁺ and Vip⁺ / Nms⁺ populations, similar to the enrichment of eGFP in the corresponding clusters from *Per1-eGFP* mice. Venus expression was also detected in subsets of Vip⁺/Grp⁺ and Cck⁺/C1ql3⁺ neurons, but it was not identified as a marker gene for these clusters and its overall levels were much weaker (Fig. 5D, E). This broader expression of Venus than eGFP likely reflects differences between the transgenes^22,24^

We next examined gene expression over time within the Venus-enriched Avp*⁺ /* Nms*⁺* and Vip*⁺ /* Nms*⁺* populations. Comparing ZT2 with ZT6 identified 51 genes in *Avp*⁺ / *Nms*⁺ neurons and 44 genes in *Vip*⁺ / *Nms*⁺ neurons were upregulated at ZT2 by ≥1.4-fold, p ≤ 0.05. *Pde10a* was among the top ZT2-enriched genes in both Venus⁺ populations, and it temporal expression matching the pattern observed in Per1-eGFP mice (Fig. 5F).

Across the time course, *Pde10a* expression was highest at ZT2 and lowest at ZT14, with a ZT2/ZT14 expression ratio of 18.42 in Avp⁺/Nms⁺ neurons and 17.83 in Vip⁺ / Nms⁺ neurons. *Pde10a* expression was also detected in Venus⁻ neurons, but with much lower expression than Venus⁺ neurons at the beginning of the day.

### Pde10a-dependent cAMP regulation of membrane excitability in Venus+ SCN neurons

Next, we tested whether papaverine depolarizes Venus⁺ neurons as we had observed in eGFP⁺ SCN neurons. We performed whole-cell current-clamp recordings between ZT2 and ZT5.75, when both Venus⁺ and Venus⁻ neurons are spontaneously active. Neurons were identified by fluorescence, and baseline RMP and SFR were recorded prior to bath applying 50 μM papaverine. The data in Supplementary Fig. 5A-C show that papaverine depolarized the RMP of Venus⁺ neurons and reduced their firing frequency, shifting them into a depolarized, electrically silent state resembling that typically observed between ZT6 and ZT10 (Fig. 1C,D). In contrast, papaverine had minimal effect on Venus⁻ neurons, with little changes in their RMP or firing rate (Supplementary Fig. 5A-C). Representative current-clamp traces (Supplementary Fig. 5C) showed that papaverine depolarized Venus⁺ neurons and affected spontaneous firing, whereas unlabeled Venus⁻ neurons retained a largely similar membrane potential and firing pattern. These results show that phosphodiesterase activity selectively regulates membrane excitability in Venus⁺ SCN neurons, consistent with higher *Pde10a* expression at ZT2 than ZT6.

### cAMP levels differentially regulate excitability in SCN neurons

The differential expression of *Pde10a* between ZT2 and ZT6 only in Venus+ neurons, and the specific effect of the phosphodiesterase inhibitor papaverine on the excitability of Venus+ neurons suggested that intracellular cAMP levels are important for altering the excitability of Venus+ SCN neurons. To test this, we used 8-Br-cAMP, a membrane-permeable cAMP analog that is widely used to activate cAMP/PKA signaling as it cannot be hydrolyzed by PDEs^34,35^. 8-Br-cAMP can induce time-of-day-dependent phase shifts in neuronal activity rhythms in the SCN and modulate glutamatergic and light-induced clock resetting^34,36^, but it had not previously been tested for a role in generating the silent depolarized state of SCN neurons.

We bath applied 500 µM 8-Br-cAMP to acute SCN slices from Per1-Venus mice collected between ZT2 and ZT5. We found that applying 8-Br-cAMP depolarized the RMP of Venus⁺ neurons and reduced SFR within 4-6 minutes. In contrast, Venus⁻ neurons were largely unaffected (Fig. 6A-C). These findings demonstrate that Venus⁺ neurons are more sensitive to elevated intracellular cAMP levels than Venus⁻ neurons. The data also support the idea that cAMP signaling differentially regulates the excitability of SCN cell types. The effects of 8-Br-cAMP closely resemble those produced by papaverine, which inhibits PDE activity and should thus also elevate intracellular cAMP.

**Figure 6.**
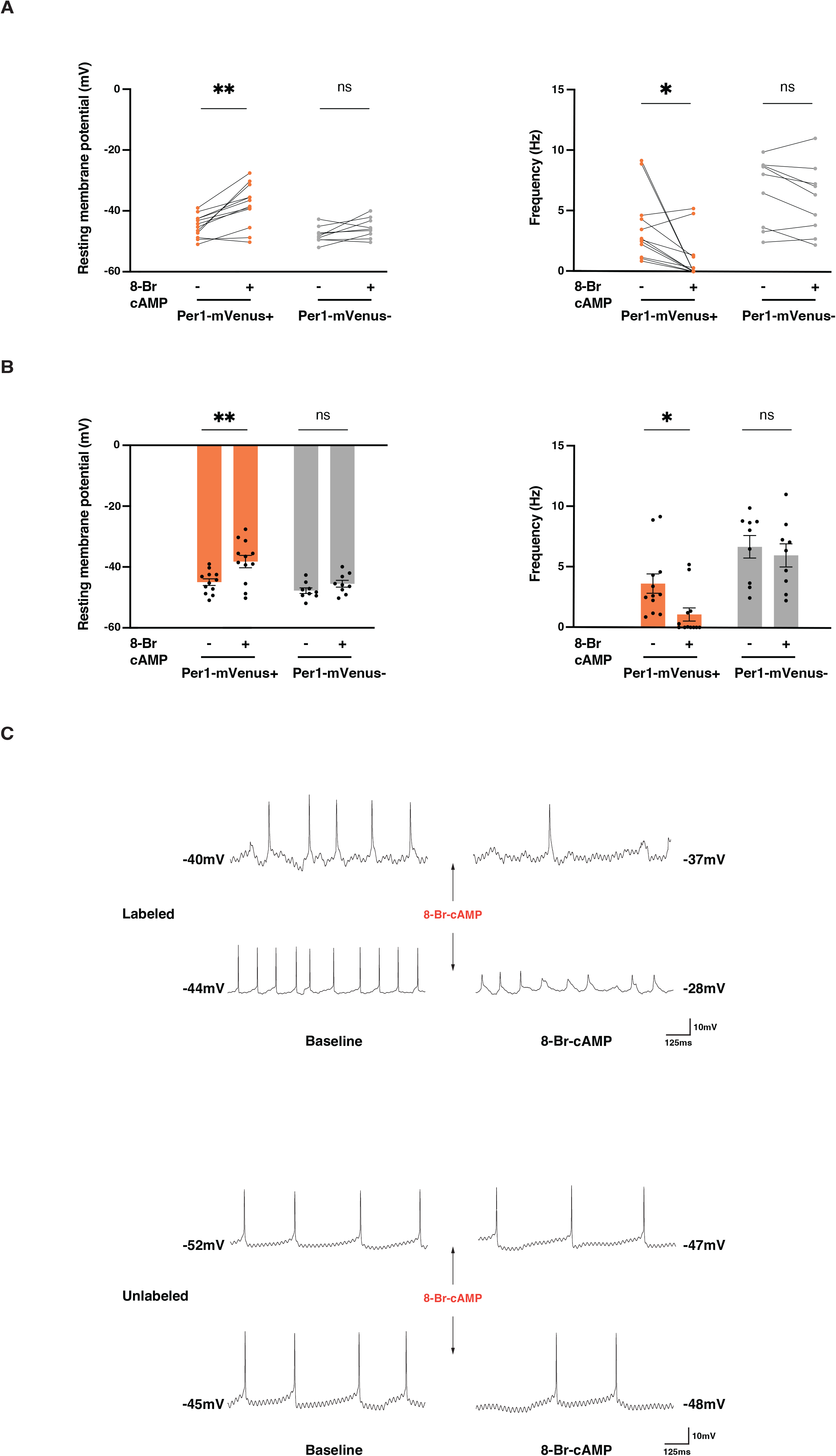
8-Br-cAMP selectively depolarize and silence Venus^+^ SCN neurons. (A) RMP (mV; left) and SFR (Hz; right) of Venus⁺ neurons and Venus⁻ neurons before and after 500 µM 8-Br-cAMP application. 8-Br-cAMP depolarized Venus⁺ neurons (paired two-tailed t-test, p = 0.0011, n = 12) but had no significant effect in Venus⁻ neurons (p = 0.0860, n = 9). Similarly, 8-Br-cAMP reduced firing in Venus⁺ neurons (p = 0.0212) but not in Venus⁻ neurons (p = 0.1112). Each line represents one neuron. (B) Quantification of RMP (mV; left) and SFR (Hz; right) before (−) and after (+) 8-Br-cAMP treatment in Venus⁺ neurons and Venus⁻ neurons. Bars represent mean ± s.e.m., and dots indicate individual neurons. Firing frequency: *p < 0.05; RMP: **p < 0.01 (paired two-tailed t-test). (C) Representative current-clamp recordings from Venus⁺ and Venus⁻ neurons before and after applying 500 µM of 8-Br-cAMP. Baseline activity was recorded after stabilization of the whole-cell configuration, and 8-Br-cAMP responses were recorded after approximately 4 min of perfusion. Arrows indicate the application of 500 µM 8-Br-cAMP. Scale bars: 10 mV, 125 ms.

Although Pde10a can hydrolyze both cAMP and cGMP^37^, prior work indicates that cAMP signaling acts mainly during the subjective day and cGMP signaling during the subjective night^34,38^. The same effect on Venus+ neuron excitability of inhibiting Pde10a activity and increasing cAMP levels is consistent with Pde10a acting as a cAMP PDE in the early morning specifically in Venus+ neurons.

Together our findings support a model in which rhythmic and high level *Pde10a* expression constrains cAMP levels in Venus⁺ neurons, thereby contributing to cell-specific regulation of SCN neuronal excitability.

## Discussion

There are two electrophysiologically distinct populations of SCN neurons^6^. One subtype exhibits gradual changes in RMP over the day, whereas a second subtype – marked by *Per1-eGFP* expression – depolarizes so strongly that they stop firing during the second half of the day. We extended this observation to a second *Per1* reporter line, and used these transgenes to identify candidate genes that underlie this switch in firing. The *Per1* reporter-positive neurons are largely Avp^+^ / Nms^+^ and Vip^+^ / Nms^+^ SCN neurons. We found that *Pde10a* expression is higher in these neurons than in the rest of the SCN, and that *Pde10a* RNA levels decrease between ZT2 and ZT6. We also found that inhibiting phosphodiesterase activity or elevating intracellular cAMP prematurely switches these Nms+ neurons into a depolarized silent state, indicating that cAMP levels normally regulate the excitability of this subset of SCN neurons – likely normally regulated by Pde10a.

Previous SCN single-cell data show that *Pde10a* RNA levels are rhythmic in Avp⁺/Nms⁺ neurons in constant darkness (DD)^26^. However, we found that *Pde10a* RNA levels are rhythmic in both Avp⁺/Nms⁺ and Vip⁺/Nms⁺ neurons in LD. This suggests that *Pde10a* expression is partly light-regulated. Indeed, Xu et al. found that *Pde10a* RNA levels are increased by a 1 hour light pulse at CT17 in Avp-, Vip- and Cck-expressing SCN neurons. In a separate study, Komal et al. found that *Pde10a* null mutant mice show increased CREB phosphorylation by light. These *Pde10a* mutant mice can also shift the phase of their locomotor activity when exposed to light during the subjective day, which is not seen in wild type mice^26,28,39^. Together with our electrophysiological findings, these studies support the idea that rhythmic *Pde10a* expression modulates cAMP signaling in defined SCN neuron subsets, shaping their time-dependent physiology.

Although papaverine can inhibit other PDEs, our transcriptomic analyses identified *Pde10a* as the most highly expressed and strongly regulated phosphodiesterase in Avp^+^ / Nms^+^ and Vip^+^ / Nms^+^ eGFP^+^ neurons. Other PDE transcripts were detected at substantially lower levels; among them, *Pde4d* showed relatively modest expression and temporal variation (Supplementary Fig. 3). Moreover, 8-Br-cAMP recapitulated the electrophysiological effects of inhibiting Pde10a, showing that elevated cAMP levels are sufficient to drive the transition towards a depolarized silent state.

We do not yet know the targets of cAMP signaling that lead to the depolarized silent state of Per1-reporter+ SCN neurons. cAMP can directly bind and regulate ion channels such as HCN^40^. cAMP can also signal through PKA and Epac in SCN neurons^21,41^ to presumably phosphorylate many substrates. Potential targets include the BK and SK potassium channels as blocking both these channels in the early day rapidly selectively switched Per1-eGFP+ SCN neurons into their afternoon depolarized silent state, with little effect on non-Per1 neurons^6^. BK channel activity is sensitive to phosphorylation, although the direction of the effect depends on the splice variant^42–44^. SK channel Ca²⁺ sensitivity and plasma-membrane availability can also be regulated by phosphorylation^45,46^

Although there is likely more than one way for a neuron to become depolarized and silent, the simplest model places Pde10a and cAMP signaling upstream of this change in conductance. We speculate that the endogenous decline in *Pde10a* RNA levels – and thus in Pde10a activity – permits cAMP-dependent phosphorylation of BK and/or SK rather than regulating ion channel mRNA levels, which are not obvious in our datasets. Future work will aim to identify the cAMP-regulated targets that link Pde10a signaling to depolarized silencing in Per1-reporter+ SCN neurons, although this will be challenging given the heterogeneity of the SCN.

Our findings add to the evidence that small molecules play central roles in circadian clocks. Cyclic nucleotides and Ca^2+^ signaling are recognized as active components of circadian timing networks rather than simple downstream outputs of the molecular clock^21,34,47,48^. Other studies have implicated redox cycles^49^ and the P-loop NTPase RUVBL2 ^50^in circadian clocks, although it is less clear how these interact with the canonical TTFL.

Outside the SCN*, Pde10* is highly expressed in striatal medium spiny neurons, where it regulates PKA signaling^51^. It will be interesting to test if inhibiting Pde10a can acutely regulate the firing of these striatal neurons by switching them to a depolarized state. Indeed, depolarized silent states are not unique to the SCN. Similar highly depolarized, silent neurons have been reported in cerebellar nuclei, and in the lateral and medial habenula^16–18^. Depolarization-induced silence in diverse brain regions suggests this is a conserved and functionally important neuronal state. Our identification of a mechanism that can govern entry into this state may therefore provide insights into the temporal control of neuronal excitability.

## Methods

### Ethics statement

All animal procedures were approved by the New York University Abu Dhabi Institutional Animal Care and Use Committee (IACUC protocol 22-000-2A3) and were performed in accordance with the NIH Guide for the Care and Use of Laboratory Animals. All researchers completed and passed the Collaborative Institutional Training Initiative Animal Care and Use core training, demonstrating familiarity with humane-care standards referenced by the USDA and the Office of Laboratory Animal Welfare.

### Mouse strains and housing conditions

Transgenic Per1 reporter mice were used for single-cell RNA-sequencing experiments. Male mice heterozygous for the mPeriod1-Venus reporter were obtained from the Jackson Laboratory (Bar Harbor, ME, USA; MMRRC stock no. 032820; FVB-Tg(Per1-Venus)33Obr/Mmjax; RRID: MMRRC_032820-JAX) and were also generously provided by Mino Belle (University of Manchester). These mice were crossed with wild-type C57BL/6J mice until a sufficient cohort of male Per1-Venus homozygotes was generated. Per1-d2eGFP mice were obtained as a generous gift from Doug McMahon (Vanderbilt University) and were bred in house. Reporter status was confirmed by genotyping. Animals were maintained in a pathogen-free facility in exhaust-ventilated, closed-system cage racks. The animal room was maintained at 21 ± 2 °C and 50 ± 10% humidity under a 12:12 LD cycle. Food and water were available ad libitum. Mice were housed with same-sex littermates until experimental assignment. At 7-8 weeks of age, male mice were transferred to individual cages and genotyped for the GFP or Venus allele. Only male mice were used for single-cell RNA-sequencing experiments.

### SCN dissection and single-cell dissociation

Mice were maintained in 12:12 LD cycles for 2–3 weeks in light- and sound-attenuated activity chambers. Mice were randomly assigned to experimental groups for tissue collection and SCN tissue was collected from light-entrained mice at specific timepoints. 4 mice were sacrificed at each timepoint and the SCN from all 4 mice was pooled to generate one biological sample for single-cell RNA-sequencing.

Animals were transcardially perfused for 1 min with 20 ml ice-cold oxygenated artificial cerebrospinal fluid (aCSF; 128 mM NaCl, 3 mM KCl, 1.25 mM NaH₂PO₄, 10 mM D-glucose, 24 mM NaHCO₃, 2 mM CaCl₂ and 2 mM MgCl₂; pH 7.4, 295-305 mOsm). 250 µm thick coronal brain slices containing the SCN were sectioned using a DTK-1000 microslicer in sucrose-substituted aCSF, with NaCl replaced by 254 mM sucrose. Bilateral SCN punches were collected and enzymatically dissociated using a Worthington Papain dissociation kit (LK003150) as described previously (Moffitt et al., 2019). Briefly, tissue punches were incubated for 45 min at 37 °C in papain buffer containing 8 U ml⁻¹ papain, 100 U ml⁻¹ DNase I, 0.8 mM kynurenic acid (Sigma, K3375), 1× GlutaMAX (Gibco, 35050-061), 0.05 mM 2-Amino-5 phosphonopentanoic acid (APV; Sigma, A5282), 0.01 mM Y-27632 dihydrochloride (Sigma, Y0503), 0.2X B27 supplement (Gibco, 17504001) and 1% D (+)-trehalose dihydrate (Sigma, T9531) in Hibernate A (Gibco, A1247501). Cells were gently triturated, centrifuged, and resuspended in 0.04% BSA (Sigma, A7906) made in 1X DPBS lacking calcium and magnesium (Gibco 14190-144). The resulting single-cell suspension was filtered through a 20 µm pluriStrainer (pluriSelect, 43-10020-40) and adjusted to approximately 400 cells / µl.

### Custom mouse reference genomes

Custom mouse transcriptome references were generated to enable quantification of reporter-derived transcripts. The *Mus musculus* GRCm38.98 reference genome was modified to include either d2eGFP or mVenus reporter gene sequences. For the former, the d2eGFP sequence was concatenated to the reference FASTA file, and the following annotation entry was appended to the corresponding GTF file:

d2EGFPensemblexon1846.+.transcript_id “eGFP”; gene_id “eGFP”; gene_name “eGFP”; exon_number “1”;

The modified eGFP-containing reference was indexed using Cell Ranger. For the mVenus reference, the mVenus sequence corresponding to NCBI accession DQ092360.1 was concatenated to the reference FASTA file, and the following annotation entry was appended to the GTF file:

DQ092360.1ensemblexon1720.+.transcript_id “mVenus”; gene_id “mVenus”; gene_name “mVenus”;

The modified mVenus-containing reference was also indexed using Cell Ranger.

### Single cell RNA sequencing (scRNA-Seq) analysis

Illumina sequencing reads were aligned to the mouse reference transcriptome using the 10x Genomics Cell Ranger pipeline. Gene-barcode matrices were imported into Seurat, and individual libraries were merged with sample-specific cell identifiers. Cells were retained if they contained 1,000–35,000 UMIs, 800–6,000 detected genes, and ≤10% mitochondrial transcripts. Genes not detected in any retained cell were removed. Mitochondrial and ribosomal genes were identified and excluded from variable-feature sets used for downstream integration and clustering. Filtered datasets were processed in Seurat. Counts from each replicate were normalized and variance-stabilized using SCTransform, regressing out total UMI count, detected gene number, mitochondrial percentage, and ribosomal percentage. Replicates were first integrated within each time point using SCT-based integration, 3,000 variable features, and the first 30 CCA dimensions. Integrated time point objects were then integrated across experimental conditions using FindIntegrationAnchors and IntegrateData. The final integrated object was scaled and subjected to principal component analysis. The first 30 principal components were used for shared-nearest-neighbor graph construction, Louvain clustering, t-SNE, and UMAP visualization. Clustering was performed across multiple resolutions, and selected neuronal clusters were further subsetted based on known SCN and neuropeptidergic marker expression.

The neuronal subset was reprocessed by recalculating variable features, excluding mitochondrial and ribosomal genes, and repeating PCA, UMAP, t-SNE, neighbor finding, and clustering. Differential expression analysis was performed using Seurat’s FindMarkers function with the Wilcoxon rank-sum test. For neuronal analysis, cell identities were defined by combining neuronal subcluster identity with time point, enabling within-cluster time point comparisons and selected group-wise contrasts. p values were adjusted for multiple testing, and genes with adjusted p value ≤ 0.05 and log fold-change > 0.25 were considered differentially expressed.

### Rhythmic gene analysis

Rhythmic gene analysis was performed primarily using pseudobulk cosinor regression. Raw single-cell counts were summed within each biological replicate, ZT time point, and model comparison group. Pseudobulk groups with fewer than 10 cells were excluded. Pseudobulk counts were converted to log2(CPM + 1), and genes with mean CPM < 1 across pseudobulk samples were excluded from the primary OLS analysis. ZT was encoded with 24 h cosine and sine terms. For each gene, three nested OLS models were fit: a null model with group only, a shared-rhythm model with group plus cosine and sine terms, and a full model with group, cosine/sine terms, and group-by-cosine and group-by-sine interactions. Rhythmicity was tested by comparing the full model against the null model, and differential rhythmicity by comparing the full model against the shared-rhythm model. P-values were adjusted using the Benjamini-Hochberg procedure. Mesor, amplitude, peak phase, amplitude differences, and circular phase differences were derived from the fitted coefficients.

As a mixed-model validation layer, the same pseudobulk count exports were analyzed with edgeR, limma/voom, and DREAM. Genes were filtered with edgeR filterByExpr, TMM-normalized, transformed with voomWithDreamWeights, and modeled with 24 h cosine/sine terms, group effects, and group-by-rhythm interactions. When at least three replicate-by-ZT blocks were available, the model included a random intercept for batch_id, defined as the interaction of biological replicate and ZT time point; otherwise, a fixed-effect model without the random intercept was fit. Rhythmicity was assessed using moderated joint tests on all rhythm-related coefficients in the full model, and differential rhythmicity using moderated joint tests on the group-by-cosine and group-by-sine interaction terms. Benjamini-Hochberg FDR correction was applied to the DREAM rhythmicity, differential rhythmicity, and mesor-difference tests. Mesor, amplitude, peak phase, and circular phase differences were computed from the DREAM fitted coefficients.

For selected clock and clock-associated genes, mesor, amplitude, and phase differences were additionally assessed using CircaCompare on log2(CPM + 1) pseudobulk expression with a 24 h period. CircaCompare was run separately for each two-group contrast. P-values for rhythmicity, mesor difference, amplitude difference, and phase difference were adjusted using Benjamini-Hochberg correction within each contrast and p-value family across the targeted gene panel.

### Brain-slice preparation for electrophysiology

Male mice at least 4 weeks of age were maintained in a 12:12 LD cycle and used for electrophysiological recordings. Male mice were used for all electrophysiological recordings, and were either heterozygous or homozygous for Per1-GFP or Per1-Venus. Before dissection, sucrose-substituted artificial cerebrospinal fluid (sucrose aCSF) and recording aCSF were equilibrated with 95% O₂ and 5% CO₂ for at least 30 min on ice. Sucrose aCSF and dissection instruments were further chilled until the solution reached a slushy consistency. Mice were deeply anaesthetized with isoflurane (0.2 ml, Vet-USP), transcardially perfused with ice-cold carbogenated aCSF and rapidly decapitated. Brains were quickly removed and transferred to chilled sucrose aCSF containing the following: 254 mM sucrose, 25 mM NaHCO_3_, 1.25 mM NaH_2_PO_4_, 10 mM glucose, 3 mM KCl, 2 mM MgSO_4_ and 2 mM CaCl_2_; pH 7.2–7.4, 290-310 mOsm. After approximately 5 min in chilled sucrose aCSF, brains were trimmed while preserving the optic chiasm as an anatomical landmark for the SCN. Coronal hypothalamic slices containing the SCN were prepared at 250 µm thickness using a Ted Pella DTK-1000 tissue slicer. Slices were then transferred to carbogenated recording aCSF containing the following: 128 mM NaCl, 25 mM NaHCO₃, 1.25 mM NaH₂PO₄, 10 mM glucose, 3 mM KCl, 2 mM MgCl₂ or 2 mM MgSO₄ and 2 mM CaCl₂; pH 7.2–7.4, 290–310 mOsm. Slices were incubated at 32 °C for 30-45 min and then maintained at room temperature for at least 30 min before transfer to the recording chamber.

### Whole-cell patch-clamp electrophysiology

SCN-containing slices were transferred to an RC-26 recording chamber (Warner Instruments) mounted on the stage of a fixed-stage upright Olympus microscope equipped with a water-immersion objective and a Hamamatsu ORCA-Spark digital CMOS camera with LED light source. Slices were continuously superfused at 2.5 ml min⁻¹ with aCSF equilibrated with 95% O₂ and 5% CO₂. Recordings were performed at 27–32 °C. Whole-cell patch-clamp recordings were performed on visually identified fluorescent and non-fluorescent neurons within the SCN.

Patch electrodes were pulled from borosilicate glass capillaries (World Precision Instruments) using a P-97 puller (Sutter Instruments). Recording pipettes had resistances of 4-8 MΩ when filled with internal solution. The standard internal solution contained the following: 115 mM K-gluconate, 20 mM KCl, 1.5 mM MgCl_2_, 0.1 mM EGTA, 10 mM HEPES, 2 mM MgATP, 0.5 mM GTP and 10 mM phosphocreatine. The pH was adjusted to 7.25-7.30, and osmolality to 280-300 mOsm. Recordings were performed using a Multiclamp 700B amplifier, and data were acquired using a Digidata 1550B digitizer and Clampfit 10.5 software (Molecular Devices, California, USA). Cells were approached while maintaining slight positive pressure. After formation of a high-resistance seal, typically up to 16 GΩ, negative pressure was applied. A second pulse of negative pressure, together with the amplifier zap function, was used to rupture the membrane and establish the whole-cell configuration. The standard extracellular solution for all recordings was aCSF.

Pharmacological agents were dissolved in MgCl₂-containing aCSF and bath-applied using a peristaltic pump-driven perfusion system. Access resistance was monitored throughout the recording, and cells were excluded from further analysis if access resistance changed by more than 40%. Series resistance was monitored during the experiments and was typically 10-25 MΩ. Data were analyzed using Clampfit 10.7.

### Drug application for electrophysiology

For pharmacological experiments, aCSF was supplemented with either 50 µM papaverine or 8-Br-cAMP 500 µM. Drugs were bath-applied continuously by perfusion. Resting membrane potential was recorded after membrane break-in and stabilization of the whole-cell configuration. Drug effects were quantified 4 mins after the start of drug perfusion. For all drug recordings, MgSO_4_-containing aCSF was replaced with MgCl_2_-containing aCSF. All other components were the same as those in the recording aCSF described above except that 2 mM MgCl_2_ replaced 2 mM MgSO_4_.

### Current injection

To verify that drug-induced changes in firing were not due to technical failure or loss of cell viability, cells were tested by injecting hyperpolarizing current up to −20 pA. Most cells responded to hyperpolarizing current injection with rebound firing. In rare cases in which the drug effect was strong and washout or current injection could not be assessed, cell-health parameters, including membrane capacitance and access resistance before and after drug application, were examined. Cells were included in the analysis only if recording parameters remained stable and within an acceptable range throughout the recording, and were excluded if access resistance changed by more than 40%.

### Statistical analysis

Statistical analysis was performed in R studio v1.4.1717 or Prism 11 with data presented as mean ± s.e.m. For electrophysiological recordings, individual data points represent single neurons, and paired lines indicate repeated measurements from the same neuron before and after current injection or pharmacological manipulation. Comparisons between reporter-positive and reporter-negative neurons recorded within the same Zeitgeber-time interval were performed using unpaired two-tailed t-tests with Welch’s correction. Within cell comparisons before and after hyperpolarizing current injection, papaverine treatment or 8-Br-cAMP treatment were performed using paired two-tailed t-tests.

For all analysis, statistical significance was defined as p < 0.05. Significance levels are indicated in the figures as follows: ns, not significant; *p < 0.05; **p < 0.01; ***p < 0.001; ****p < 0.0001.

## Contributions

J.B. and D.C. conceived the study. P.N. designed, planned and optimized the experiments and performed the electrophysiology and single-cell RNA-sequencing experiments. M.Š. and S.K. assisted with the single-cell RNA-sequencing workflow. P.N., M.Š. and S.K. analyzed the single-cell data. S.K. assisted with figures. G.A.S performed rhythmicity analysis. P.N. interpreted the data. P.N., J.B., D.C. wrote the manuscript. J.B. and D.C. supervised the study, acquired funding.

## Acknowledgements

We thank Doug McMahon and Mino Belle for *Per1-eGFP* and Per1-Venus mice respectively. We also thank Hugh Piggins and Mino Belle for advice on electrophysiology. This work was supported by Tamkeen via an NYU Abu Dhabi (NYUAD) Research Institute Award to the NYUAD Center for Genomics and Systems Biology. Cell sorting was performed at the NYUAD CGSB FACS Core and Bioinformatics assistance was provided by the NYUAD CGSB Bioinformatics Core. We thank Marc Arnoux for assistance with deep sequencing and technical support, and Mehar Sultana for assistance with deep sequencing and technical support for the FACS experiments. We also thank Dr. Rachid Rezgui (NYUAD Microscopy Core Technology Platform) for help with imaging to confirm GFP expression, and Nizar Drou (CGSB Bioinformatics Core) for generating the genomes for *Per1-eGFP* and *Per1-Venus* mice.

## Supplementary figure legends

**Supplementary Figure 1:**
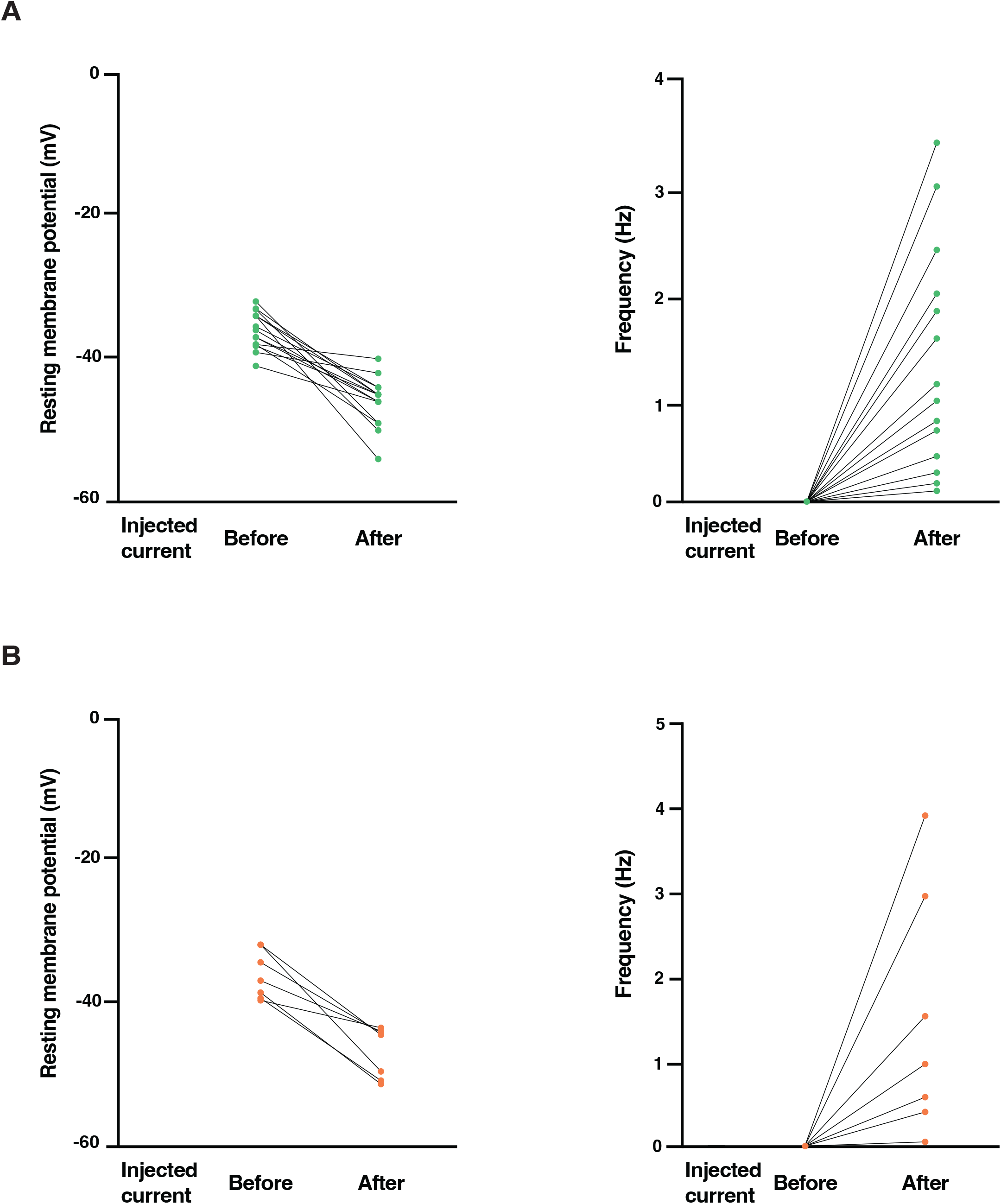
Paired whole-cell recordings from individual eGFP⁺ and Venus⁺ neurons showing the effect of hyperpolarizing current injection on RMP (left) and SFR (right). (A) Injecting a current of up to −20pA repolarized eGFP⁺ neurons (-40 mV to -54 mV) and restored their firing (0.1-3.1 Hz). Lines connect paired measurements from the same neuron before (left) and after (right) current injection. (B) Injecting a current of up to −20pA repolarized Venus⁺ neurons (-43.6 mV to -51.6 mV) and restored their firing (0.05-3.9 Hz). Lines connect paired measurements from the same neuron before (left) and after (right) current injection.

**Supplementary Figure 2.**
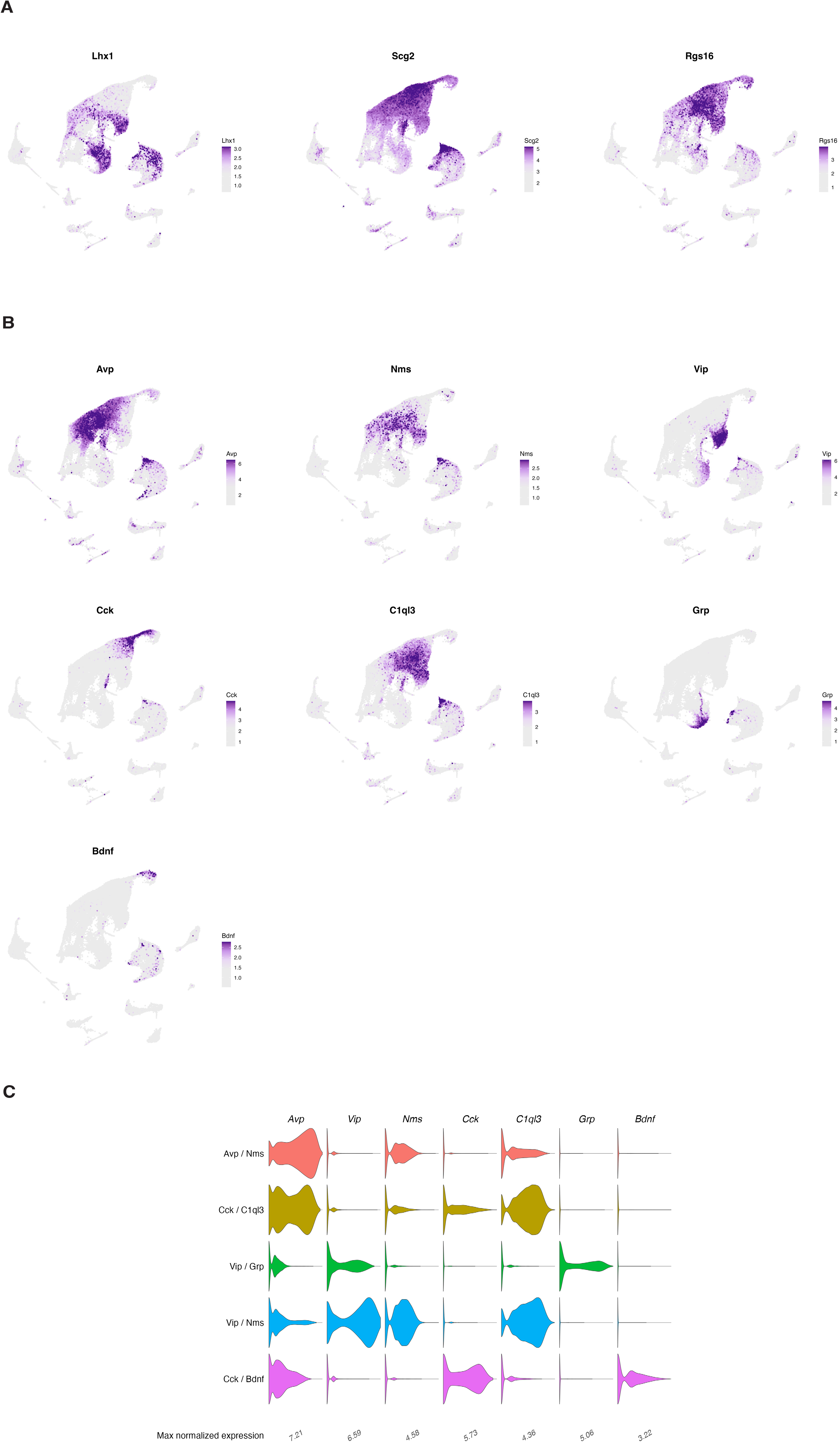
Expression of marker genes used to define SCN neuronal subtypes in Per1-eGFP mice. (A) UMAP feature plots showing expression of the SCN-associated genes *Lhx1*, *Scg2* and *Rgs16* across SCN clusters. Color intensity indicates normalized gene expression, from low (grey) to high (purple). These genes are highly expressed in SCN neurons relative to surrounding cell populations. (B) UMAP feature plots showing expression of the neuropeptide and subtype marker genes *Avp*, *Nms*, *Vip*, *Cck*, *C1ql3*, *Grp* and *Bdnf*. Color intensity indicates normalized gene expression. (C) Violin plots showing the distribution of these marker genes across the five transcriptionally defined SCN neuronal subtypes: *Avp*⁺/*Nms*⁺, *Cck*⁺/*C1ql3*⁺, *Vip*⁺/*Grp*⁺, *Vip*⁺/*Nms*⁺ and *Cck*⁺/*Bdnf*⁺. Violin width reflects the distribution of expression values within each subtype.

**Supplementary Figure 3.**
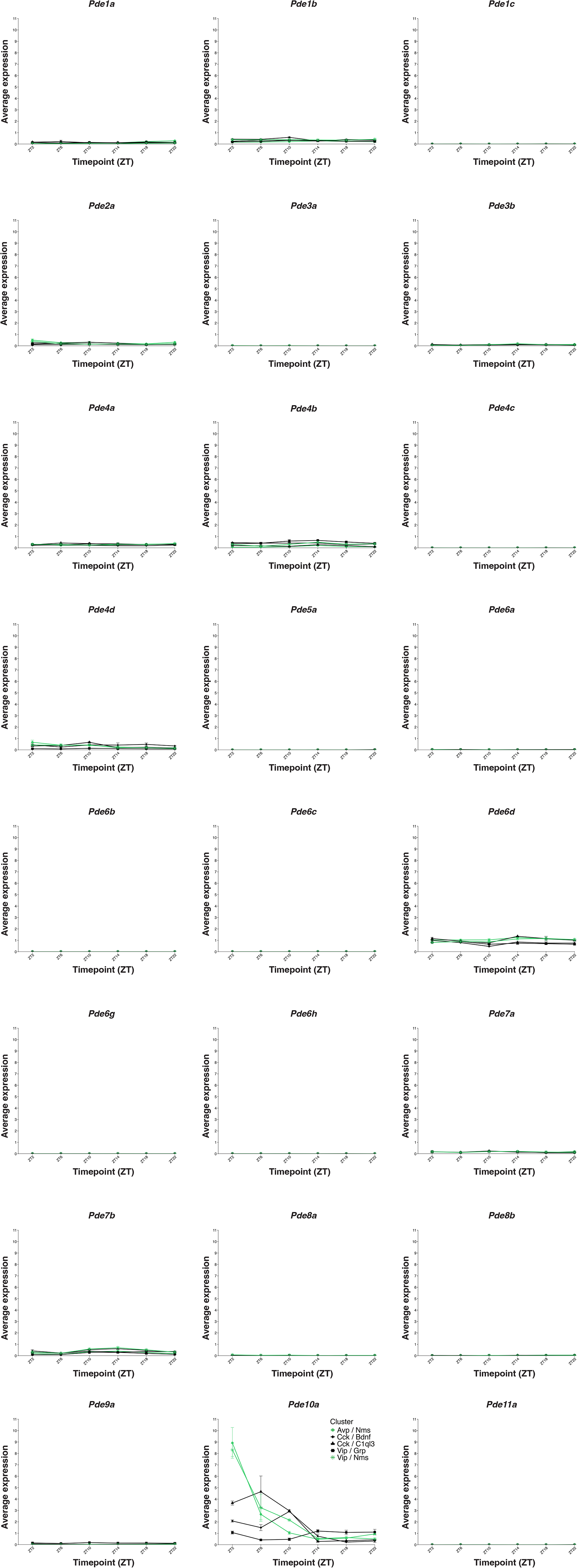
Temporal expression of phosphodiesterase genes across SCN neuronal subtypes. Line plots show average normalized RNA expression of 24 *Pde* genes across six Zeitgeber time points (ZT2, ZT6, ZT10, ZT14, ZT18 and ZT22) in the five transcriptionally defined SCN neuronal subtypes: *Avp*⁺/*Nms*⁺, *Cck*⁺/*Bdnf*⁺, *Cck*⁺/*C1ql3*⁺, *Vip*⁺/*Grp*⁺ and *Vip*⁺/*Nms*⁺. Each line represents one neuronal subtype, and points show mean expression with error bars indicating s.e.m. All genes are plotted using the same y-axis scale to allow direct comparison across the *Pde* family. Most *Pde* transcripts showed low expression across subtypes, whereas *Pde10a* displayed time-dependent pattern, with higher early-day expression in *Avp*⁺/*Nms*⁺ and *Vip*⁺/*Nms*⁺ neurons and later profiles in the *Cck*⁺ clusters.

**Supplementary Figure 4.**
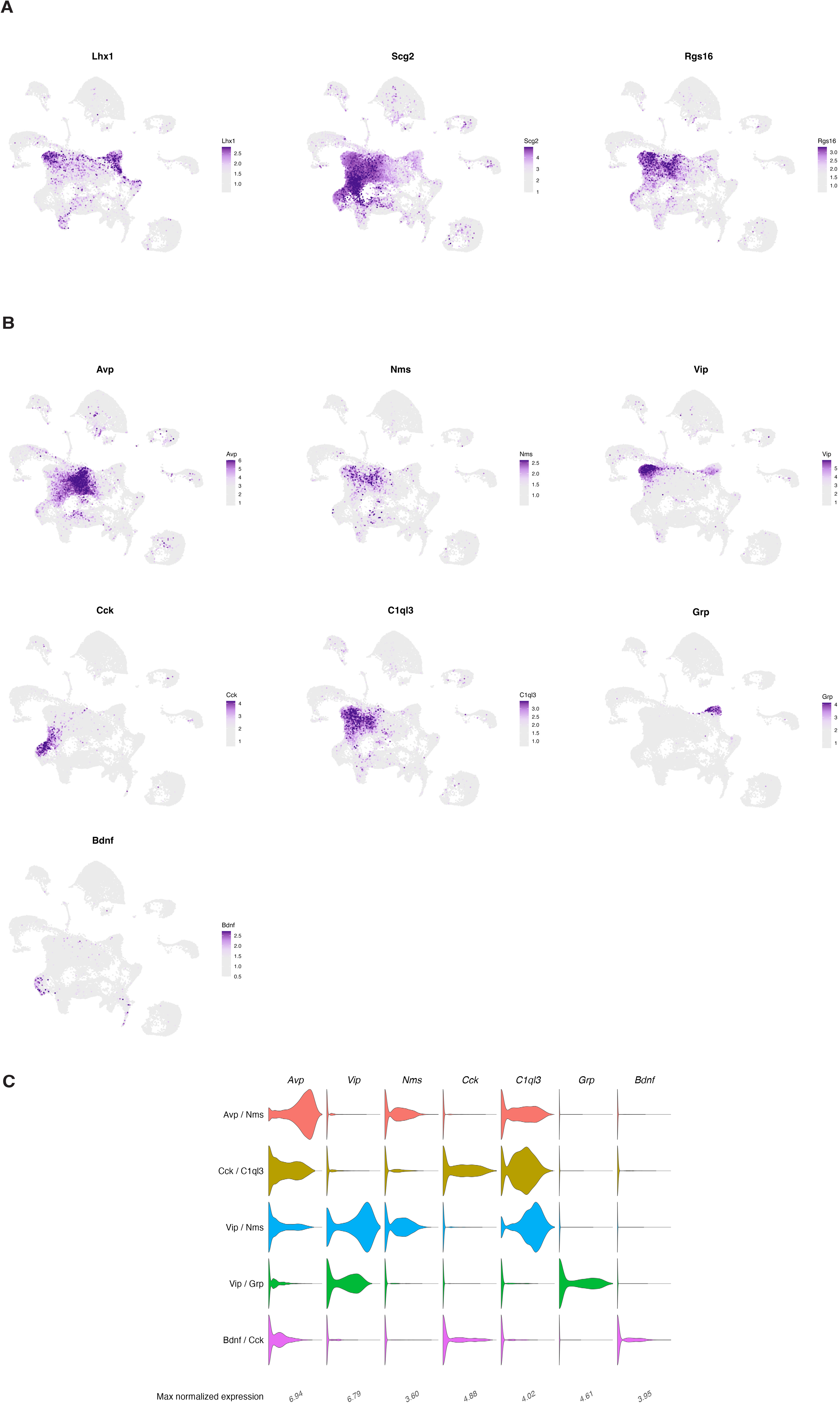
Expression of marker genes used to define SCN neuronal subtypes in Per1-Venus mice. (A) UMAP feature plots showing expression of the SCN-associated genes Lhx1, Scg2 and Rgs16 across SCN clusters from Per1-Venus mice. Color intensity indicates normalized gene expression, from low (grey) to high (purple). These genes are highly expressed in SCN neurons relative to surrounding cell populations. (B) UMAP feature plots showing expression of the neuropeptide and subtype marker genes Avp, Nms, Vip, Cck, C1ql3, Grp and Bdnf. Color intensity indicates normalized gene expression. (C) Violin plots showing the distribution of these marker genes across the five transcriptionally defined SCN neuronal subtypes: Avp⁺/Nms⁺, Cck⁺/C1ql3⁺, Vip⁺/Nms⁺, Vip⁺/Grp⁺ and Cck⁺/ Bdnf⁺. Violin width reflects the distribution of expression values within each subtype.

**Supplementary Figure 5.**
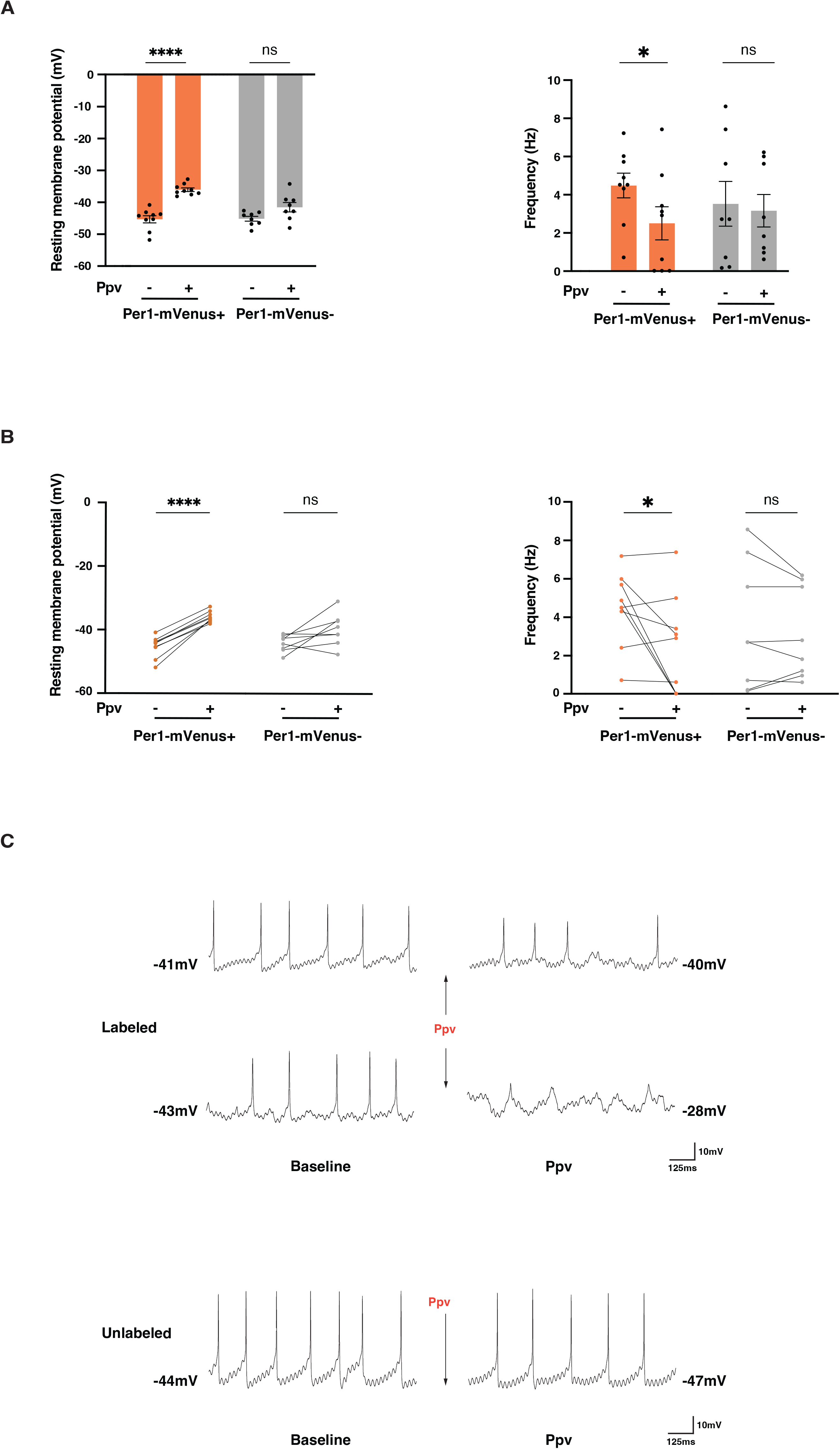
Effects of papaverine on membrane excitability in Per1-Venus SCN neurons. (A) RMP (mV, left) and SFR (Hz, right) of Per1-Venus⁺ and neighbouring Venus⁻ SCN neurons recorded between ZT2 and ZT5.75 before and after bath application of papaverine hydrochloride (Ppv; 50 µM). Ppv depolarized Venus⁺ neurons (paired two-tailed t-test, p < 0.0001, n = 9) but did not significantly alter RMP in Venus⁻ neurons (*p* = 0.061, *n* = 8). Ppv also reduced SFR in Venus⁺ neurons (*p* = 0.0472, n= 9), but not in Venus⁻ neurons (p *=* 0.3996, n = 8). Bars show mean ± s.e.m; dots represent individual neurons. \**p* < 0.05, \*\*\*\**p* < 0.0001; ns, not significant. (B) Paired RMP (mV; left) and SFR (Hz; right) measurements before and after Ppv treatment. Lines connect measurements obtained from the same neuron. Statistical comparisons and sample sizes are as in A. *p<0.05; ****p<0.0001; ns, not significant. (C) Representative current-clamp recordings from Per1-Venus⁺ and Per1-Venus⁻ SCN neurons under baseline conditions and following Ppv application. Per1-Venus⁺ neurons displayed two distinct responses to Ppv: a reduction in firing frequency and the membrane potential oscillations. In Per1-Venus⁻ neurons, the effect of Ppv was less pronounced, with no marked change in firing activity. Baseline activity was recorded 1 min after establishing the whole-cell configuration, and Ppv responses were recorded after 4 min of perfusion. Arrows indicate Ppv application. Scale bars: 10 mV, 125 ms.

